# Kilometre-scale jaguar swimming reveals permeable hydropower barriers: implications for conservation in the Cerrado hotspot

**DOI:** 10.1101/2025.09.05.674446

**Authors:** Leandro Silveira, Giselle Bastos Alves, Anah Tereza de Almeida Jácomo, Tiago Jácomo, Fabio Hudson Souza Soares, Gabriel Caputo de Carvalho, Lucas Gonçalves da Silva

**Author notes:** **Corresponding Authors** Leandro Silveira, Lucas Gonçalves da Silva.

## Abstract

Hydropower reservoirs are expanding rapidly across tropical biomes, yet their function as barriers or filters to carnivore movement remains poorly understood. Here, we report the first confirmed long-distance swim by a jaguar (*Panthera onca*) across an artificial lake and discuss its implications for landscape connectivity. Camera traps around Serra da Mesa Reservoir (Central Brazil; 1,784 km^2^; 54.4 km^3^) photographed an adult male on the mainland and later on a forested island. Flank-pattern matching confirmed a 100% identity between records. Geodesic analysis identified two possible routes: a direct 2.48 km crossing or an alternative path involving a stepping-stone islet (1.06 km + 1.27 km). In the absence of evidence for use of the islet, we conservatively adopt the largest continuous water segment—1.27 km—as the minimum distance swum. This represents nearly six times the longest previously verified jaguar swim (≈ 200 m). The event supports a graduated-resistance framework in which water bodies exceeding 1 km impose high, but not insurmountable, costs to movement. We propose an ordinal aquatic-cost scale (1 = < 300 m; 3 = 300–1,000 m with stepping-stones; 6 = > 1,000 m open water) for use in circuit-based connectivity models. By contrasting this record with predator collapse on > 1 km islands in Lake Guri and with genetic segregation across the Amazon River, we demonstrate that aquatic permeability is strongly context-dependent. Incorporating rare but consequential swims into environmental impact assessments will improve corridor design and better inform hydropower licensing throughout the jaguar’s range.

**Highligts:**

– Extends the longest recorded jaguar swim by almost six times.
– Demonstrates that reservoirs act as costly filters rather than absolute barriers for large carnivores.
– Introduces an ordinal aquatic-cost scale to enhance connectivity models.
– Informs hydropower impact assessments and the design of amphibious corridors for mammal conservation.

## 1. Introduction

The jaguar (*Panthera onca*) is the largest felid in the Americas and an apex predator whose presence regulates trophic interactions and maintains key ecological processes in the Neotropics (Terborgh, 1992; Estes et al., 2011). Historically distributed from the southwestern United States to northern Argentina, the species has lost approximately 50% of its original range due to habitat conversion, fragmentation, and direct persecution, resulting in severe population declines and multiple regional extinctions (Sanderson et al., 2002; de la Torre et al., 2017). Like other large carnivores, jaguars depend on dispersal to sustain gene flow, avoid inbreeding and recolonize vacant habitats (Rabinowitz & Zeller, 2010). However, landscape-genetic modeling indicates that when functional connectivity falls below critical thresholds (e.g., ∼30% of natural cover), populations may experience sharp reductions in genetic diversity within just one generation, jeopardizing long-term viability (Pfeifer et al., 2017; Chapron et al., 2023). Mapping jaguar movement corridors across increasingly human-dominated landscapes is thus a central challenge for conservation ecology, requiring integrative approaches that combine camera-trap data, telemetry, and predictive connectivity models (Morato et al., 2016; Cushman et al., 2018).

Traditionally, jaguars are closely associated with forested habitats adjacent to permanent water bodies (Sunquist and Sunquist, 2014) and are recognized as highly capable swimmers (Rabelo et al., 2019). However, the extent to which large rivers or wide reservoirs constrain (or alternatively facilitate) long-distance dispersal remains poorly understood. Hydropower development alone has already inundated more than 25,400 km^2^ of jaguar habitat, with 429 additional projects planned, effectively transforming continuous forest into archipelagos of isolated islands (Palmeirim and Gibson, 2021). The Guri Reservoir in Venezuela is an classic example that illustrates the ecological consequences of such fragmentation: after inundation, islands separated by >1 km became inaccessible to terrestrial predators, leading to a trophic collapse within just three decades (Terborgh et al., 2001). Despite these impacts, landscape connectivity models from the Jaguar Corridor Initiative assign a resistance value of 100 to lake stretches exceeding 1 km, compared with 20–40 for fluvial channels shorter than 300 m, suggesting that aquatic barriers represent a gradient of permeability rather than absolute obstacles (Zeller et al., 2012).

In this context, direct observations challenge the assumption that large water bodies act as absolute barriers. African lions (*Panthera leo*) have been documented crossing over 1 km of open water in the Kazinga Channel (Braczkowski et al., 2024). Regarding jaguars, the longest anecdotal distance previously recorded is ca. 200 m (Holt, 1932). These rare events suggest that, under favorable conditions (e.g., warm water, low currents, presence of stepping-stone islands) large felids may occasionally exploit aquatic corridors that appear to be initially insurmountable (Elbroch et al., 2022).

While reservoirs represent relatively recent barriers, the Amazon River has served as a long-standing biogeographic boundary. Analyses of 715 bp of mitochondrial DNA and 29 microsatellite loci revealed an FST of 0.34 between opposite banks, indicating substantial, but not complete, genetic segregation; occasional crossings appear to occur during low-water periods or via mid-river islands (Eizirik et al., 2001). An even more extreme case occurs on the Maracá–Jipioca islands off Amapá State, Brazil, located 6–10 km from the mainland. Camera-trap surveys and genetic sampling have documented 21 jaguars within only 149 km^2^ (density ≥ 11 ind./100 km^2^), including breeding females. Occupancy models indicate that population viability likely depends on periodic marine dispersal (Duarte et al., 2023).

Beyond such instances, jaguars exhibit remarkable ecological plasticity. They exploit fish-rich Amazonian várzea floodplains (Ramalho et al., 2021), persist in semi-arid Caatinga refugia (Astete et al., 2017), and based on GPS tracking of 116 individuals cover more than 120 km per month even under moderate human disturbance, reducing their movements only when disturbance indices surpass 2.5–5.5 (Morant et al., 2025). This behavioral flexibility indicates those long aquatic environments crossing events, while rare and energetically demanding, fall within the species’ natural behavioral repertoire.

Despite growing evidence, quantitative data on jaguars swimming distances exceeding one kilometer, particularly in human-modified landscapes, remain virtually nonexistent. Here, we analyzed a documented male jaguar crossing event in open waters from Serra da Mesa Reservoir, located in the Cerrado of Brazil, considering prepositions such as: (i) if jaguars movements collapses on islands isolated longer than 1 km (proposed by: Holt, 1932); (ii) the hypothesis of Amazon River’s acting as an incomplete barrier for jaguar individuals movement (proposed by: Eizirik et al., 2001), and; (iii) the distinct pattern represented by marine-influenced population of Maracá-Jipioca (proposed by: Duarte et al., 2023). This study aims to clarify the conditions under which water bodies function as barriers, filters, or corridors for *P. onca*. These insights are critical for refining aquatic-cost parameters in connectivity models, informing infrastructure planning, prioritizing restoration measures, and promoting conservations actions that consider the amphibious movement capabilities of one of Latin America’s flagship predators.

## 2. Methods

### 2.1. Study Area

The dispersal event was recorded in the reservoir of the Serra da Mesa Hydroelectric Dam (13°45′ S, 48°25′ W), located in northern Goiás State, Brazil, within the upper Tocantins River basin. The region is part of the Cerrado biome, a Neotropical savanna characterized by vegetation ranging from open grasslands to dense cerradões and gallery forests along permanent streams (Ribeiro & Walter, 2008). The local topography consists of a dissected plateau incised by deep valleys and mesas. The impoundment of the dam in 1996 created a highly sinuous shoreline and hundreds of islands of varying size. The reservoir currently covers 1,784 km^2^ and stores 54.4 km^3^ of water, making it the largest artificial lake in Brazil by volume (Caramaschi et al., 2012; FURNAS, 2025). The documented swim occurred across a section approximately 4 km wide, between the southern mainland margin (Minaçu municipality) and one of the largest forested islands formed following inundation.

### 2.2. Exploratory Survey

We conducted an exploratory jaguar survey initiated in April 2020 around the Serra da Mesa reservoir. Four camera-trap stations were deployed: three on the mainland and one on an island (Figure 1). Each station was equipped with a passive infrared camera (Suntek HC802A) positioned approximately 50 cm above the ground along terrestrial trails identified based on tracks, scats, and local reports. Cameras operated continuously (24h day^−1^) at high sensitivity, with a minimum 30-second interval between captures, and were checked every 30–40 days to replace batteries and memory cards. Although each station’s single camera captures only one flank, high-quality images allow individual identification through spot-pattern analysis (Silveira et al., 2003).

**Figure 1.**
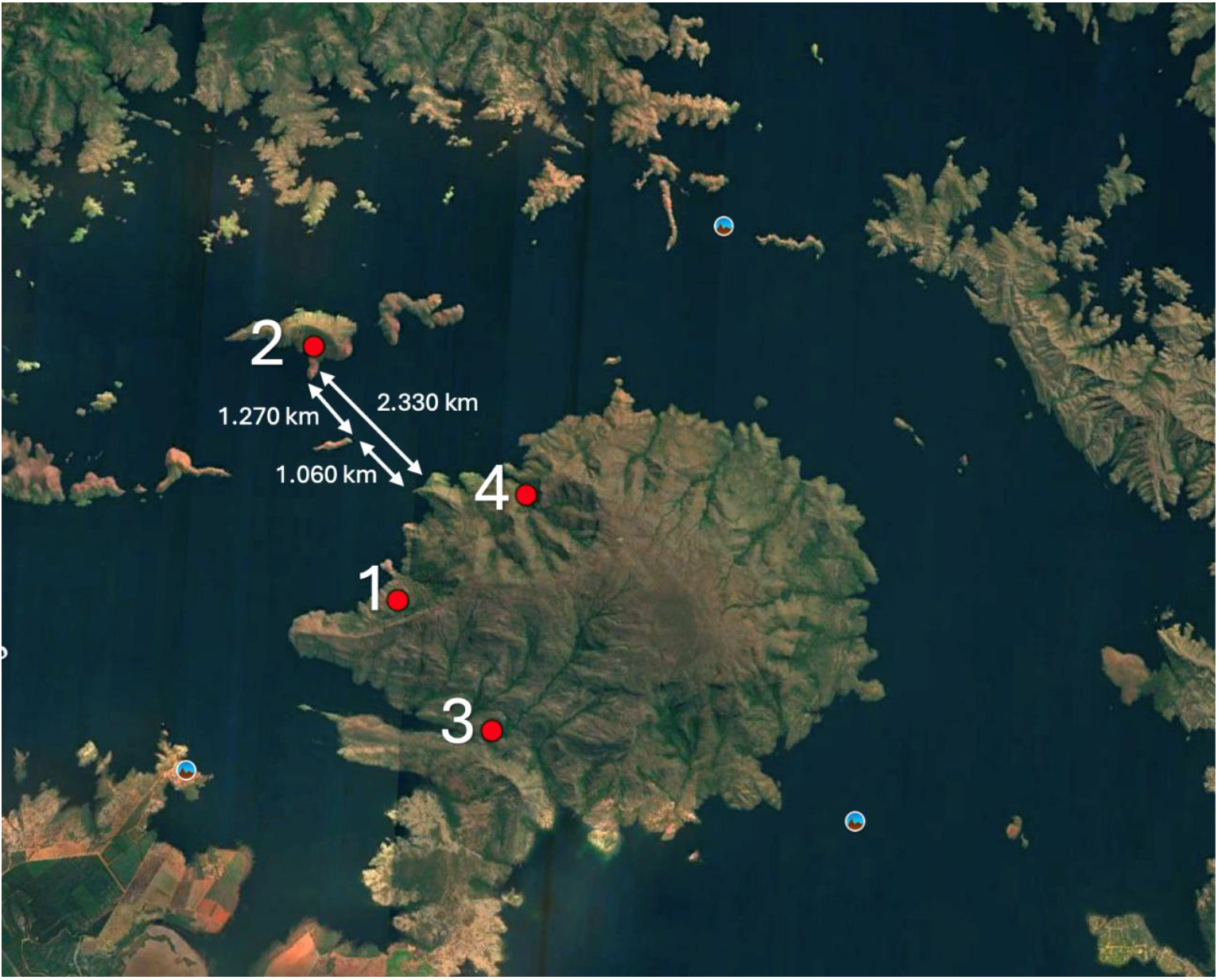
Study area and camera-trap locations around Serra da Mesa Reservoir. Camera trap locations: 1 – first jaguar record; 2 – second record; 3 and 4 no records.

**Figure 2.**
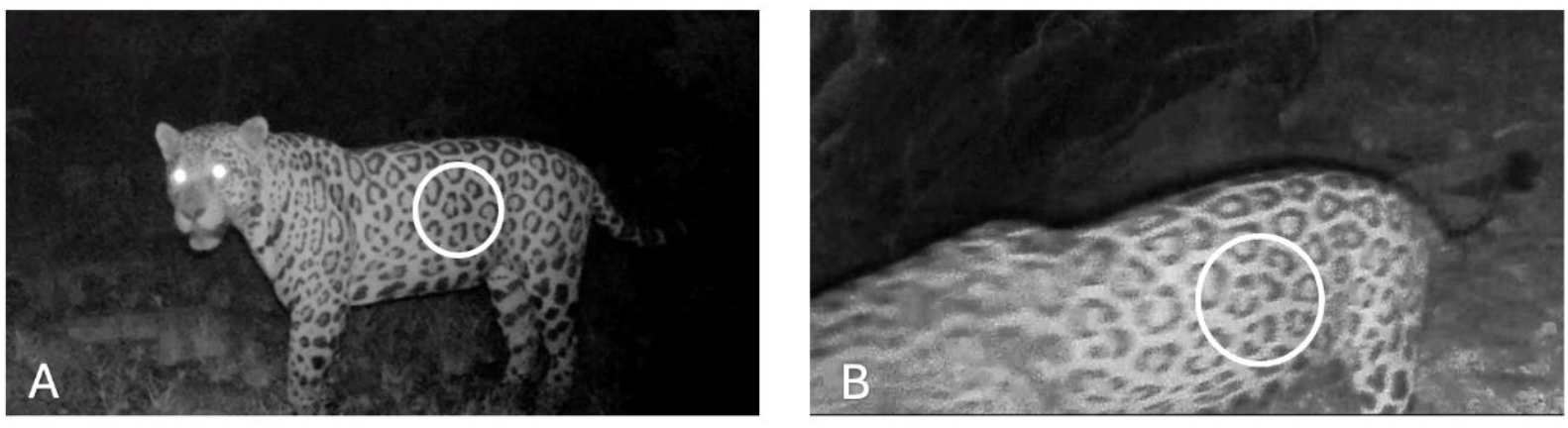
Camera-trap records in mainland (A) and island (B).

## 3. Results

Between April 2020 and March 2025, four camera-trap stations were systematically deployed in the reservoir area of the Serra da Mesa Hydroelectric Power Dam, northern Goiás State, Brazil. On May 2nd, 2020, an adult male jaguar (*Panthera onca*) was photographed for the first time on the northern shore of the reservoir (14°10′ S, 48°17′ W). Rosette-pattern analysis confirmed a 100% match across all subsequent records, allowing confident individual identification. Based on the authors’ field experience, the animal displayed robust adult morphology, with a visually estimated body mass exceeding 80 kg, which indicates a minimum age of ≥ 3 years.

After more than four years of monitoring, this same individual was recorded again on August 8th, 2024, by a camera trap installed on Island X (14°09′ S, 48°14′ W), a large landmass situated within the flooded area of the reservoir. To estimate the minimum swimming distance required for this dispersal event, we calculated straight-line trajectories in QGIS 3.36, using 30 m resolution SRTM topographic data, and evaluated two scenarios (Figure 1): (i) a direct crossing between the mainland and Island X, totaling 2.48 km; (ii) a stepwise route via an intermediate islet (Islet Y), consisting of 1.06 km (mainland → Islet Y) plus 1.27 km (Islet Y → Island X), for a total of 2.33 km.

Since the exact departure point on the mainland could not be determined, we adopted the conservative estimate based on the shortest continuous aquatic stretch that the jaguar would necessarily have swum—1.27 km (Islet Y → Island X), nearly six times the longest anecdotal distance previously recorded (ca. 200 m; Holt, 1932). This represents the longest uninterrupted segment of open water crossing required to reach the island where the record was obtained. Given the scarcity of documented long-distance swims by large terrestrial carnivores, this event provides a rare quantitative measure of jaguar aquatic dispersal capacity within a human-modified landscape.

## 4. Discussion

### 4.1. Historic Background

Early naturalists reported felids swimming across rivers up to ∼200 m wide, suggesting that water functions as a relative rather than absolute barrier (Holt, 1932). Modern GPS telemetry corroborates this view: jaguars in fragmented landscapes move >120 km month^−1^ (≈4 km day^−1^), demonstrating high dispersal capacity and potential for extended aquatic movements (Morato et al., 2016). The 1.27 km crossing documented here expands the verified swimming range of jaguars nearly six-fold and constitutes the longest measured open-water movement for Panthera onca to date. Taken together, these observations warrant a reassessment of the species’ dispersal potential across large South American waterbodies.

### 4.2. Ecological Plasticity

Jaguars exhibit remarkable behavioural versatility. They hunt semi-aquatic prey in the Pantanal (Quigley & Crawshaw, 1992), persist in the xeric Caatinga through montane refugia and limited water sources (Astete et al., 2017), and adopt an amphibious lifestyle during Amazonian flood pulses (Ramalho et al., 2021). The 1.27 km reservoir crossing indicates that this plasticity also encompasses uninterrupted open-water expanses. Comparable kilometre-scale swims by African lions across Uganda’s Kazinga Channel (Braczkowski et al., 2024) confirm that large carnivores can negotiate substantial aquatic gaps when ecological incentives are strong. Collectively, these lines of evidence demonstrate that carnivore movement models must explicitly incorporate amphibious capacity.

### 4.3. Reservoir Permeability

Connectivity models often assign near-absolute resistance to hydropower reservoirs (Venter et al., 2016). Empirical evidence from Lake Guri, where large predators failed to recolonize islands >1 km from the mainland (Terborgh et al., 2001), seems to support that assumption. However, our record from Serra da Mesa suggests that reservoir resistance may be graduated rather than binary. In this case, warm water, low boat traffic, and a stepping-stone islet apparently lowered the functional cost sufficiently for an adult male jaguar to swim 1.27 km of open water. We therefore propose a provisional three-tier aquatic-cost scale for jaguars: (1) Low cost (ordinal value 1): ≤300 m when at least one islet is available; (2) Medium cost (value 3): 300–1,000 m with an islet, or; (3) High cost (value 6): >1,000 m with no islets.

These dimensionless values are hypotheses that require calibration with GPS-telemetry and camera-trap data collected under diverse hydroclimatic conditions. They provide a first step toward incorporating water-crossing energetics into regional connectivity models. At broader spatial scales, landscape-level modeling shows that hydropower reservoirs can severely curtail connectivity for wide-ranging carnivores when treated as impermeable barriers (Palmeirim & Gibson, 2021).

Published telemetry studies seldom report open-water swims exceeding a few hundred metres, and no peer-reviewed record exists of a jaguar swimming >1 km. The most comprehensive dataset to date—117 GPS-collared jaguars across five Neotropical countries (Morato et al., 2018)—did not document any single aquatic crossing above that threshold. Consequently, our 1.27 km observation represents the longest confirmed swim for the species, albeit from a single individual. Field surveys confirmed the absence of bridges, culverts, or emergent land corridors linking Island X to the mainland during the 2020–2025 dry seasons, reinforcing the inference that the jaguar crossed the reservoir aquatically.

### 4.4. Management and conservation implications

Because some reservoirs are at least partially permeable, conservation planning should consider amphibious corridors that integrate riparian habitat with aquatic stepping-stones. Maintaining gentle shore gradients, preserving riparian vegetation, and safeguarding islets ≤1 km apart can reduce swimming costs for jaguars and other taxa. Evidence of primates swimming tens of metres between Amazonian islands (e.g. Pavelka et al., 2025) highlights potential multi-taxon benefits. Incorporating such corridor elements can help sustain trophic integrity in reservoir-dominated landscapes.

## 5. Conclusions

Our documentation of a 1.27 km swim by a jaguar across the Serra da Mesa Reservoir challenges the prevailing assumption that large water bodies function as absolute barriers to carnivore movement. This record—nearly six times longer than any previously reported jaguar swim—reveals a far greater capacity for aquatic dispersal than previously recognized. By introducing an ordinal aquatic-cost scale, we provide a framework for integrating graduated resistance values into connectivity models, advancing beyond simplistic binary barrier assumptions. These insights have direct relevance for hydropower impact assessments and corridor planning, highlighting that the strategic retention of riparian habitat and placement of stepping-stone islets can sustain landscape permeability for wide-ranging carnivores. Future research should aim to quantify the frequency of such crossings and test the applicability of the proposed cost scale across multiple reservoir systems, thereby supporting evidence-based conservation strategies in hydropower-dominated landscapes.

## 6. Acknowledgements

Authors would like thank to Jaguar Conservation Fund for financial assistance.

## 7. Data Availability Statement

Camera-trap photographs, GIS layers, and analysis scripts will be deposited in Zenodo (DOI to be provided) upon article acceptance; meanwhile, they are available from the corresponding author upon reasonable request.

## 9. CRediT authorship contribution statement

**Leandro Silveira**: Conceptualization; Methodology, Writing–original draft; Writing–review and editing; Supervision. **Giselle Bastos Alves:** Writing–original draft; Writing–review and editing. **Anah Tereza de Almeida Jácomo:** Funding acquisition; Writing–review and editing; Supervision. **Tiago Jácomo:** Methodology; Investigation; Writing–original draft; Writing–review and editing. **Fabio Hudson Souza Soares:** Investigation; Formal analysis; Writing–review and editing. **Gabriel Caputo de Carvalho:** Investigation; Formal analysis; Writing–review and editing. **Lucas Gonçalves da Silva:** Formal analysis; Writing–review and editing.

